# Circulating Microvesicle-Associated Inducible Nitric Oxide Synthase Is a Novel Therapeutic Target to Treat Sepsis: Current Status and Future Considerations

**DOI:** 10.1101/2021.11.08.467816

**Authors:** Robert J Webber, Richard M Sweet, Douglas S Webber

**Affiliations:** Research & Diagnostic Antibodies, Las Vegas, NV, USA; School of Medicine, University of California San Francisco and Renal Department, Zuckerberg San Francisco General Hospital, San Francisco, CA, USA

**Keywords:** sepsis, therapeutic target, inducible nitric oxide synthase, iNOS, microvesicles, nanoparticles

## Abstract

**Objective:** To determine if mitigating the harmful effects of circulating microvesicle-associated inducible nitric oxide (MV-A iNOS) *in vivo* increases the survival of challenged mice in three different mouse models of sepsis.

**Design:** The ability of anti-MV-A iNOS monoclonal antibodies (mAbs) to rescue challenged mice was assessed using three different mouse models of sepsis.

**Setting:** The vivarium of a research laboratory

**Subjects:** Balb/c mice

Interventions: Mice were challenged with an LD_80_ dose of either lipopolysaccharide (LPS / endotoxin), TNFα, or MV-A iNOS and then treated at various times after the challenge with saline as control or with an anti-MV-A iNOS mAb as a potential immunotherapeutic to treat sepsis.

**Measurement and Main Results:** Each group of mice was checked daily for survivors, and Kaplan-Meier survival curves were constructed. Five different murine anti-MV-A iNOS mAbs from our panel of 24 murine anti-MV-A iNOS mAbs (1) were found to rescue some of the challenged mice. All five murine mAbs were used to genetically engineer humanized anti-MV-A iNOS mAbs by inserting the murine complementarity-determining regions (CDRs) into a human IgG_1,kappa_ scaffold and expressing the humanized mAbs in CHO cells. Three humanized anti-MV-A iNOS mAbs were effective at rescuing mice from sepsis in three different animal models of sepsis. The effectiveness of the treatment was both time and dose dependent. Humanized anti-MV-A iNOS rHJ mAb could rescue up to 80% of the challenged animal if administered early and at a high dose.

**Conclusions:** Our conclusions are MV-A iNOS is a novel therapeutic target to treat sepsis; anti-MV-A iNOS mAbs can mitigate the harmful effects of MV-A iNOS; the neutralizing mAb’s efficacy is both time and dose dependent; and a specifically targeted immunotherapeutic for MV-A iNOS could potentially save tens-of-thousands of lives annually and could result in improved antibiotic stewardship.

## Introduction

Sepsis is an enormous medical problem globally (2). The current sepsis definition, Sepsis-3, is now defined as life-threatening organ dysfunction caused by a dysregulated host response to infection (3). Over the past several years, the need to rethink therapeutics to treat sepsis and diagnostic tests to identify individuals at a high risk of becoming septic has been recognized (4,5,6). As is widely known, more than two dozen Phase 3 clinical trials on numerous candidate therapies for sepsis have failed during the past 30+ years (4,5). At present, not a single specifically targeted drug is approved for the treatment of sepsis. Likewise, numerous candidate biomarkers have been proposed to detect sepsis, but only one, plasma inducible nitric oxide synthase (iNOS) appears to be specific for the onset of sepsis (6,7). The discovery of the normally intracellular enzyme iNOS circulating in the blood in microvesicles and its relationship to the sepsis pathology as both a specific biomarker for the onset of sepsis and a novel therapeutic target to treat sepsis (6,8) led us to develop a recombinant humanized anti-microvesicle associated (MV-A) iNOS IgG _1,kappa_ monoclonal antibody (rHJ mAb) as an efficacious candidate immunotherapeutic (8).

Recently, the number of deaths worldwide involving sepsis was estimated to be more than 11 million annually, 19.2% of all deaths each year (2). This estimate was made based upon the analysis of millions of death certificates from 2017. It has also been recently reported that almost all deaths due to the COVID-19 pandemic result from sepsis (9,10,11). The in-hospital and long-term economic and societal burden of sepsis makes it one of the most pressing patient care issues worldwide. However, despite intense research efforts and the investment of billions of dollars over the past 30+ years by pharmaceutical and biotechnology companies and by government research institutes worldwide, a targeted therapy to treat sepsis has still not been developed (4,5).

The US Department of Health and Human Services (DHHS) has estimated the direct cost of sepsis to the US healthcare system to be greater than $60 billion annually (12,13,14). Health economic analyses have estimated the direct cost of sepsis is only 28% of its total cost to society – the other 72% of its cost is due primarily to lost productivity from individuals who either die of sepsis or are disabled and are unable to provide for themselves and their families after surviving an episode of sepsis (15,16,17).

As the sepsis pathology progresses, organ dysfunction often occurs and deteriorates into multiple organ failure and death (18,19,20). Sepsis is estimated to be involved in up to 50% of all hospital deaths, and morbidity and mortality remain unacceptably high in septic patients mainly due to the lack of a specific and targeted treatment (21,22,23). Also, mortality has been reported to increase approximately 5% per year for the first five years after discharge due to the long-term health complications of sepsis (24,25).

The current definition of sepsis (3), Sepsis-3, is based upon an individual patient’s SOFA score (or quick SOFA score, qSOFA) and superseded the prior definition, now called Sepsis-2, that was based upon the SIRS criteria and a confirmed or suspected infection (26). Over the last few years, numerous research groups have demonstrated that when the Sepsis-2 and Sepsis-3 definitions are applied to actual patients enrolled in clinical studies, the two definitions define different septic patient subpopulations with only a partial overlap (27,28). Because a spectrum of symptoms exists, many septic patients do not display the same or similar symptoms. Thus, even defining what sepsis is (or is not) has been a challenge and has hindered the development of a specific therapeutic (3,4,5,18,19,26,29).

The clinical need for a specific and efficacious therapeutic to treat sepsis is widely recognized and well documented (3,4,5,18,19,21,26). Most clinicians in this field agree that a targeted therapy for the sepsis pathology would be a major medical breakthrough and would vastly improve best practices treatment and care for septic patients. An effective targeted therapeutic for sepsis should decrease mortality and morbidity and save healthcare systems worldwide billions of dollars annually (12,13,14,15,16).

For the last few years, our research team has been investigating a novel form of the normally intracellular iNOS enzyme that is not customarily found in blood. However, during the onset of sepsis, iNOS is found in circulating microvesicle (MV) nanoparticles as microvesicle-associated iNOS (MV-A iNOS) in whole blood and plasma samples (6,7). Based upon our data and data from independent investigators (30,31,32), circulating MV-A iNOS appears to play a major role in the onset of the sepsis cascade by producing toxic quantities of NO in sites distal from the site of an infection and thereby causing cellular damage and organ dysfunction. Data presented herein demonstrate that MV-A iNOS is a validated new therapeutic target for sepsis, and that anti-MV-A iNOS mAbs can effectively stop the harmful systemic effects caused by circulating MV-A iNOS. Thus, a recombinant humanized anti-MV-A iNOS mAb can be the first effective immunotherapeutic drug to treat sepsis and to meet this significant unmet medical need.

## Materials and Methods

### Radio-labeling MV-A iNOS as Tracer

To determine the half-life of MV-A iNOS in blood and if the uptake of MV-A iNOS by organs is altered by lipopolysaccharide (LPS/endotoxin), MV-A iNOS was purified by centrifugation followed by column chromatography. The fractions containing the purified MV-A iNOS were pooled, dialyzed, concentrated, and radio-iodinated using iodogen and carrier free Na^125^I. The ^125^I-labeled MV-A iNOS was used to determine the T½ of MV-A iNOS in blood and was also used in a series of pulse-chase experiments to determine its uptake by different organs.

### T½ determination for ^125^I-MV-A iNOS in Blood

Five mice were injected IV with 4 x 10^6^ dpm of ^125^I-MV-A iNOS, and at time points, a 50 μl blood sample was collected from the tail vein of each mouse in hematocrit tubes. The blood was centrifuged, and an aliquot of plasma was counted in a scintillation well gamma counter. The data was plotted, and the half-life calculated.

### Tissue Distribution of MV-A iNOS

Eight groups of three mice (N=3) were primed with a sub-lethal dose of LPS or given a saline injection as controls; 4 hours later, they were injected IV with 4 x 10^6^ dpm of ^125^I-MV-A iNOS; and 60 min later euthanized by ether anesthesia. Before dissection, each animal was subjected to transcardial whole-body perfusion with 20 ml of sterile saline to remove blood from all the organs. Heart, liver, intestines, and spinal cord were rapidly dissected, weighed, and counted for their content of ^125^I on a scintillation well gamma counter. The results for each mouse as disintegrations per minute (dpm) ^125^I per mg tissue (dpm/mg) was calculated for each organ. The average dpm/mg was then calculated for each organ for all groups.

### Genetically Engineered Recombinant Humanized Anti-MV-A iNOS Monoclonal Antibodies (mAbs)

After determining that five of the mAbs in our panel of 28 murine anti-iNOS mAbs (1) possessed *in vivo* neutralizing activity as judged by their ability to rescue challenged animals from death in three different animal models of sepsis, each was humanized by inserting the mouse CDRs into human IgG_1_ heavy chains and human kappa light chains. The humanized mAbs were stably transfected into and expressed as recombinant humanized IgG_1,kappa_ monoclonal antibody in *dhfr-* CHO cells. After a series of cloning, selection, and amplification rounds, each of the stable rCHO cell lines was cyropreserved as a master cell bank. All the recombinant humanized anti-MV-A iNOS mAb clones were tested for *in vivo* neutralization of MV-A iNOS in our mouse bioassay (8,33).

### Kaplan-Meier Survival Curves

To determine if rHJ mAb was effective as a candidate treatment for sepsis, groups of mice were injected IV with LPS at 6.0 mg/kg (an LD_80_ dose of LPS), and 0, 2, and 6 hours later were treated with either saline (control), or low dose rHJ mAb (125 ng/gm body weight), or high dose rHJ mAb (1.25 μg/gm body weight). The number of mice surviving was checked daily for 7 days, and the results were plotted as Kaplan-Meier Survival Curves. Two additional mouse models of sepsis were also utilized: one used an LD_80_ IV dose of TNFα, and the other used an LD_80_ IV dose of MV-A iNOS (8,33).

All animal studies were reviewed and approved by an Institutional Animal Care and Use Committee (IACUC).

## Results

Previously, our team reported that while conducting clinical studies on inducible nitric oxide synthase (iNOS) as a potential new blood biomarker for the onset of sepsis, it was discovered that the iNOS in blood was exclusively contained in circulating microvesicle nanoparticles and was only present in the blood of individuals who were already septic or who would become septic in the next 24 to 48 hours (6,7). A name for the circulating iNOS in blood microvesicles was created, microvesicle-associated iNOS (MV-A iNOS), since iNOS in blood appears to be present exclusively in circulating microvesicles and not as a freely soluble protein (6,7).

After discovering MV-A iNOS, experiments were performed to determine its half-life in blood and its tissue and organ distribution. Based upon the disappearance of IV administered ^125^I-MV-A iNOS from blood, the half-life of MV-A iNOS was found to be biphasic. The fast component has a T½ of 11 minutes, and the slow component has a T½ of 18 hours. To compare the changes in the distribution of MV-A iNOS to tissues and organ that resulted from a non-lethal dose of LPS, the results of the pulse-chase experiments for the heart, spinal cord, intestines, and liver were normalized to the value found for the saline treated control animals for each organ individually (Table 1). Statistically significant differences were calculated by Student’s T-test, and P values less than 0.05 are considered significant. In the heart, LPS treatment resulted in an 86% increase in the amount of ^125^I-MV-A iNOS taken-up as compared to the saline treated group. In the spinal cord, LPS treatment resulted in an 87% increase in the amount of ^125^I-MV-A iNOS taken-up as compared to the saline treated group. In the intestines, treatment with LPS resulted in a more than doubling of the amount of ^125^I-MV-A iNOS taken-up as compared to the saline treated group. However, in the liver, no discernible change was found in the amount of ^125^I-MV-A iNOS that was up taken following LPS treatment as compared to the Saline control group.

**Table 1.** Organ Uptake of ^125^I-MV-A iNOS After an LPS Challenge in Mice Normalized against Saline Controls

| Organ | Normalized DPM $^{125}\text{I}$/mg Tissue | | |
| --- | --- | --- | --- |
|  | Saline Only | Saline + LPS | P Value * |
| Heart | 100.0% | 186.4% | $P < 0.05$ |
| Spinal Cord | 100.0% | 187.1% | $P < 0.05$ |
| Intestines | 100.0% | 201.1% | $P < 0.05$ |
| Liver | 100.0% | 103.5% | No Significant Difference |

\* A difference of $P < 0.05$ is considered statistically significant

In order to determine if any of our murine anti-iNOS mAbs possesses *in vivo* neutralizing activity, our panel of 24 anti-iNOS monoclonal antibodies (1) was screened for their individual ability to neutralize *in vivo* the lethal effects of circulating MV-A iNOS in our mouse bioassay (8,33). Five of the murine anti-iNOS mAbs in our panel were found to be effective at rescuing mice from a lethal challenge of MV-A iNOS (Figure 1). All five of the *in vivo* neutralizing murine anti-MV-A iNOS mAbs were humanized by genetically engineering techniques. All the humanized anti-MV-A iNOS clones were tested for *in vivo* neutralization of MV-A iNOS.

**Figure 1:**
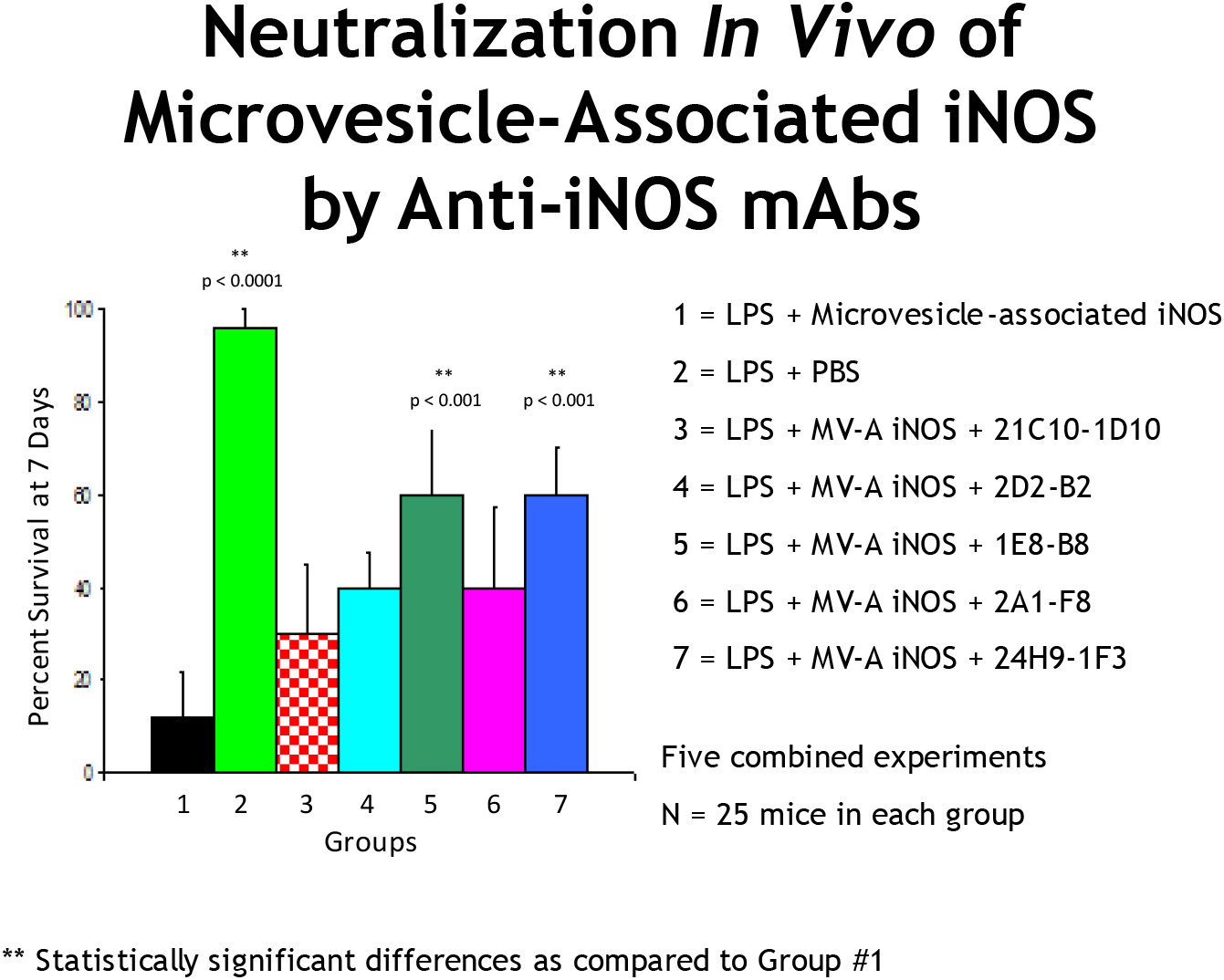
These are the combined data from five different experiments in which we tested all our murine anti-iNOS mAbs for their individual ability to neutralize *in vivo* the lethal effects of microvesicle-associate iNOS (MV-A iNOS). As depicted in lane #1, mice were dosed with LPS at 2 mg/kg at time zero, and that was followed with a dose of MV-A iNOS at 4 hours. At this dose of MV-A iNOS, 88% (22 of 25 died) of the challenged animals die of sepsis. As Lane #2 shows, if saline is injected at 4 hours instead of MV-A iNOS, then more than 95% (24 of 25 lived) of the animals lived because a very low dose of LPS was used to prime the animals. Many of the murine anti-iNOS mAbs in our panel (1) did not rescue mice from death by sepsis since they did not neutralize the lethal effects of MV-A iNOS *in vivo* even though they all bound to iNOS. However, intervention with the anti-iNOS mAbs illustrated in Lanes # 3 – 7 did have a beneficial effect since they rescued some of the challenged mice from death by sepsis by stopping the sepsis cascade in these animals. The anti-iNOS mAbs depicted in Lanes # 3 – 7 were selected for continued investigation as potential candidate therapeutics to treat the sepsis pathology. The mouse monoclonal antibody secreting hybridoma cell lines were used to construct recombinant humanized anti-MV-A iNOS mAbs. Humanized anti-MV-A iNOS J mAb (rHJ mAb) is our lead immunotherapeutic candidate to treat sepsis.

Three recombinant humanized anti-MV-A iNOS mAbs – rHA mAb, rHD mAb, and rHJ mAb – were found to rescue mice from a lethal challenge in three different mouse models of sepsis, an LPS model, a TNFα model, and our in-house MV-A iNOS model of sepsis (33). As is illustrated by the Kaplan-Meier Survival Curves (Figure 2) for the LPS mouse model using rHJ mAb intervention, three recombinant humanized anti-MV-A iNOS mAbs were shown to have some efficacy. Their effectiveness was both dose and time-after-challenge dependent – the higher the dose and the earlier administered, the more effective each humanized anti-MV-A iNOS mAb was at rescuing mice from death by sepsis. However, only rHJ mAb, could rescue up to 80% of the challenged animals from death by sepsis if administered within the first two hours after the LPS challenge.

**Figure 2:**
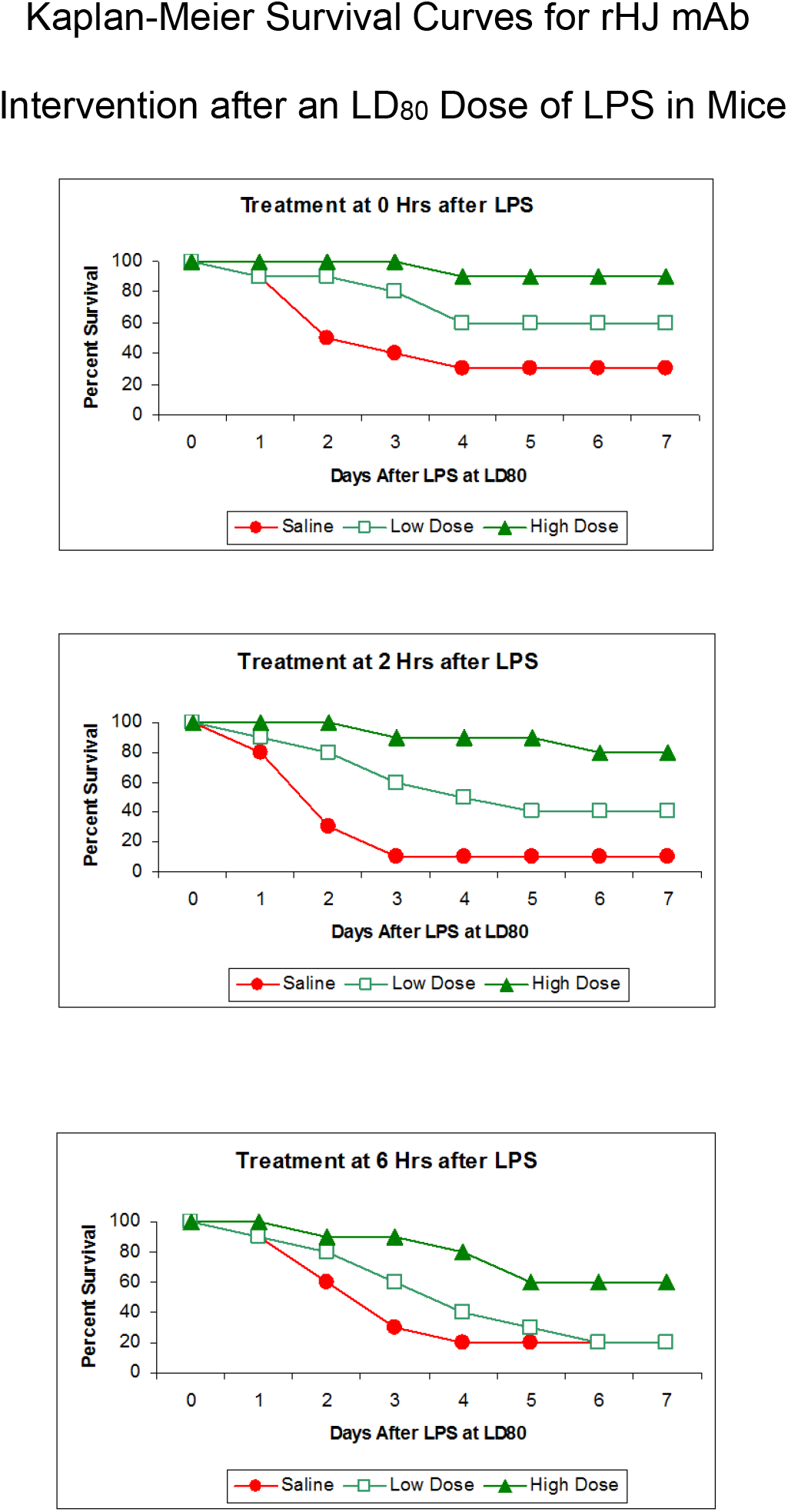
These Kaplan-Meier Survival Curves demonstrate intervention with both the Low Dose (125 ng/gm body weight) and the High Dose (1.25 μg/gm body weight) of rHJ mAb was effective at rescuing the mice from death by sepsis as compared to the Saline control group. The earlier rHJ mAb was administered and the higher the dose given, the more effective it was. Similar results were also obtained when an LD80 dose of TNFα or an LD80 dose of MV-A iNOS was given IV as the challenge.

## Discussion

Our prior data (6,7,8,33) demonstrate that what occurs in sepsis is aberrant apoptosis (Figure 3) leading to secondary necrosis of cells induced to produce iNOS, to the release into the circulatory system of microvesicle-associated iNOS, and ultimately to the life-threatening sepsis cascade. Our discovery of the release of microvesicles containing iNOS into blood has been confirmed for septic humans (30,31) and septic rats (32). After the circulating iNOS-containing microvesicles are isolated, they have been shown by us (8,33) and other investigators to cause cell damage (30), organ damage and dysfunction (31), and death (32). The left-hand side of the illustration depicts a cell that has been induced by the “inflammatory cytokine storm” to make iNOS. The red dashed line indicates that macrophages (Mø) do not recognize this cell as apoptotic: either the cell may not mark itself properly, or the macrophages may not recognize the markings properly, or there might be a local depletion of macrophages and others could not be recruited in fast enough to scavenge this induced, apoptotic cell. Thus, this apoptotic cell is not properly scavenged by a macrophage.

**Figure 3:**
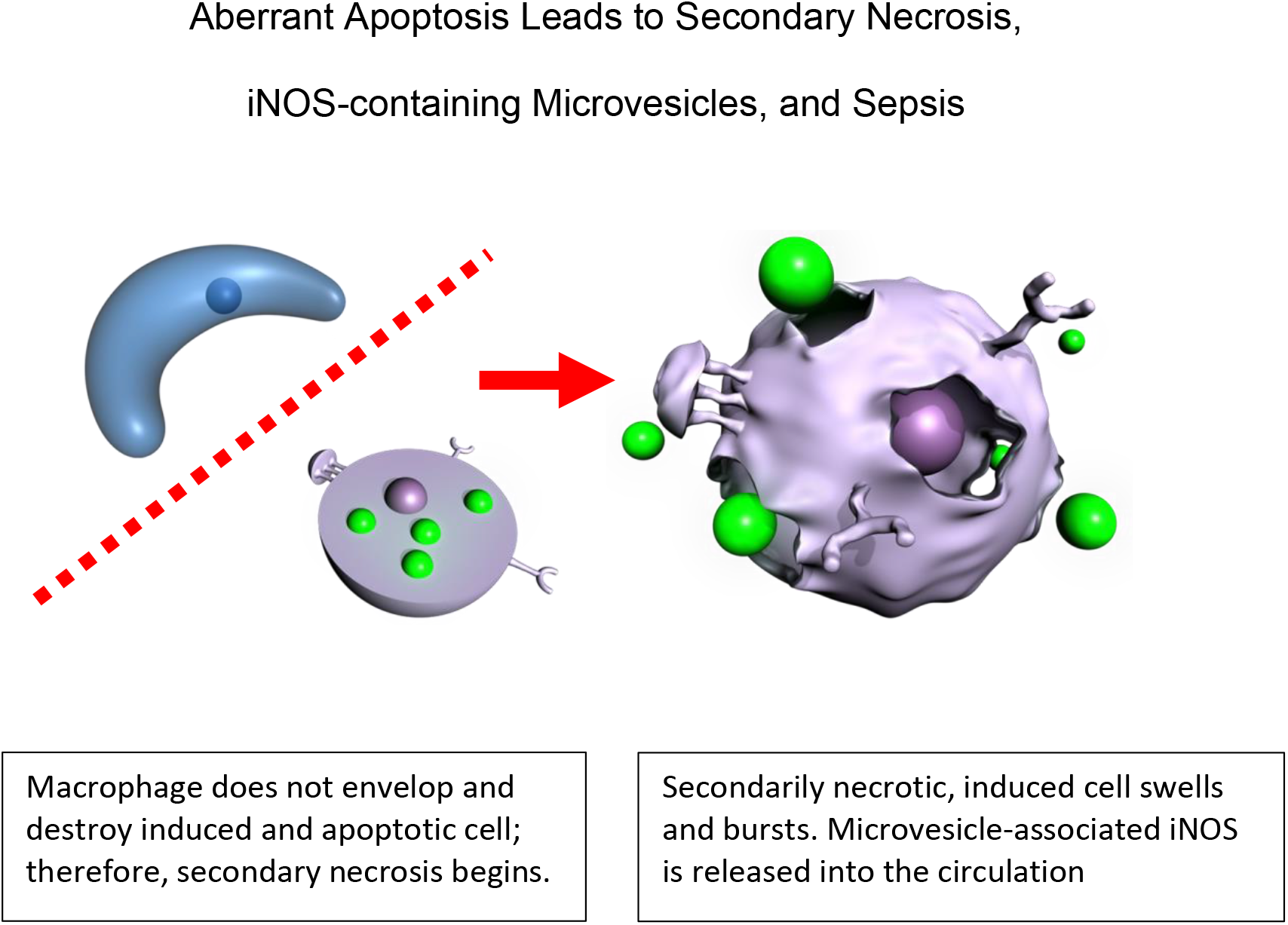
Published data by our team (6,7,8,33) demonstrate that what triggers sepsis is aberrant apoptosis which leads to secondary necrosis of cells induced to produce iNOS, to the release into the circulatory system of microvesicle-associated iNOS, and ultimately to the life-threatening sepsis cascade. This is a new pathophysiological pathway that was recently discovered and confirmed for the sepsis pathology. The left-hand side of the illustration depicts a cell that has been induced by the inflammatory cytokine storm to make iNOS. The red dashed line indicates that macrophages (Mø) do not recognize this cell as apoptotic. Thus, this induced apoptotic cell is not properly scavenged by a macrophage and instead undergoes secondary necrosis, which is depicted in the right-hand side of this illustration. The secondarily necrotic, induced cells swell, burst, and release their cellular contents into the circulatory system including iNOS-containing microvesicles (depicted as extracellular green balls).

The question is then, “What happens to this cell?” The answer is the induced, apoptotic cell undergoes secondary necrosis, which is now depicted in the right-hand side of this illustration. The secondarily necrotic, induced cells swell, burst, and release their cellular contents into the blood, and the iNOS enzyme contained in the circulating microvesicles is then delivered as cargo to receiver cells at distal sites in the body (Figure 4).

**Figure 4:**
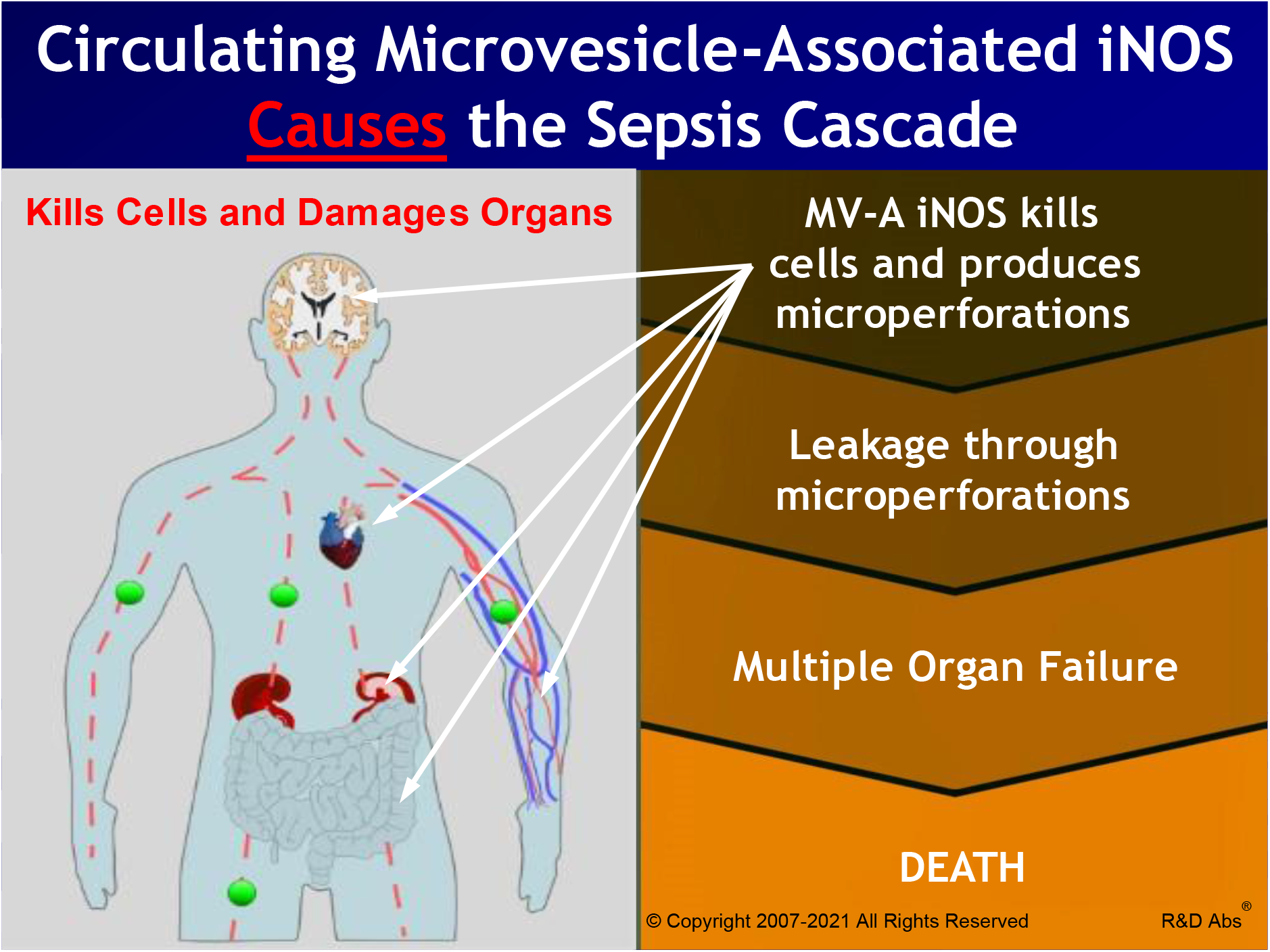
This illustration shows the causative role that microvesicle-associated iNOS (MV-A iNOS), shown as green balls, plays in sepsis. Once the MV-A iNOS is delivered as cargo contained in circulating microvesicles to receiver cells at distal sites in the body, the active iNOS enzyme is internalized into the receiver cells, where it produces toxic quantities of nitric oxide. This kills the receiver cell which produces microperforations in barrier structures that then leak, and it also damages the myocardium of the heart which leads to hemodynamic collapse and death.

While the iNOS-containing microvesicles are in the circulatory system, the iNOS is an inactive enzyme, because two of its required cofactors are not present in plasma. Our data and the data of other investigators show that these circulating microvesicles lodge onto or bind to the surface of cells and are then internalized into the receiver cells. These processes have been comprehensively described for circulating extracellular microvesicles in other human diseases (34,35,36,37). Once a microvesicle containing iNOS is intercalated into a receiver cell, the MV-A iNOS becomes an active enzyme, since it has all of its required substrates and cofactors. However, it is now a component of a cell that has never been induced, it is in an inappropriate location, and the receiver cell is out of normal cellular regulation. Once active, the iNOS enzyme produces toxic quantities of nitric oxide that results in the death of that cell. Damage to cardiomyocytes leads to the hemodynamic collapse of the heart (31). Damaged to the blood-brain barrier, the tight junctions of the intestine, the glomerular filtration units of the kidney, and the capillary beds of the circulatory system leads to leakage through these barrier structures. When a receiver cell takes up the circulating microvesicles containing iNOS at one of these locations, the internalized and active iNOS produces enough nitric oxide (NO^∸^) to kill that cell. Thus a microperforation is formed in the barrier through which leakage can occur. Microperforations in the blood-brain barrier results in plasma leaking into the brain and to neurological symptoms associated with sepsis. Microperforations in the tight junctions of the intestines leads to leakage from the lumen of the intestine into the circulatory system, and results in bacteremia and/or endotoxemia. Microperforations in the kidneys’ glomerular filtration units result in plasma proteins leaking into the urine and in renal failure. Microperforations in the capillary beds results in vascular leak syndrome including leakage into the interstitial space and an inability to maintain blood pressure. These are the classic hallmark symptoms of sepsis, severe sepsis, and septic shock caused by the circulating MV-A iNOS. This is a new pathway for sepsis that was recently discovered and confirmed.

The ability to stop the delivery of the MV-A iNOS to receiver cells with an *in vivo* neutralizing immunotherapeutic drug, such as our rHJ mAb, should be rigorously investigated because of its potential to meet this huge unmet medical need, to save lives worldwide, and to save healthcare systems billion of dollars annually.

## Conclusions

Circulating MV-A iNOS is a validated new immunotherapeutic target to treat sepsis in three different mouse models of sepsis. Kaplan-Meier Survival curves demonstrate that the effectiveness of each recombinant humanized anti-MV-A iNOS mAbs is dose and time dependent. Recombinant humanized J mAb (rHJ mAb) can rescue up to 80% of the challenged animals from death by sepsis if administered early in an episode of sepsis. Our candidate rHJ mAb immunotherapeutic to treat sepsis should rapidly be made available to treat the more than 50 million annual cases of sepsis, including septic COVID-19 patients, since almost all COVID-19 fatalities are caused by viral sepsis (9,10,11).

